# MolSpecFlow: Mass-Constrained Hybrid Flow Matching for Joint Molecular-Spectral Analysis

**DOI:** 10.64898/2026.01.28.702438

**Authors:** Yu Wang, Fan Yang, Kaikun Xu, Li Yuan, Jun Zhu, Jingjie Zhang, Zhenchao Tang, Yatao Bian, Cheng Chang, Yonghong Tian, Jianhua Yao

**Affiliations:** School of Computer Science, Peking University; School of Artificial Intelligence for Science, Peking University; AI for Life Sciences Lab, Tencent; State Key Laboratory of Medical Proteomics, National Center for Protein Sciences, Beijing; Research Unit of Proteomics-driven Cancer Precision Medicine, Chinese Academy of Medical Sciences; School of Electronic and Computer Engineering, Peking University; Tsinghua-Peking Joint Center for Life Sciences, School of Life Sciences, Tsinghua University; Department of Computer Science and Engineering, The Chinese University of Hong Kong; Department of Computer Science, National University of Singapore

## Abstract

Identifying the “dark matter” of the chemical universe requires bridging the fundamental gap between molecules and mass spectra. Existing approaches struggle to accurately map between these distinct data formats, often yielding results that are either chemically implausible or inconsistent with the physical evidence. We introduce MolSpecFlow, a unified foundation model pretrained on 100 million molecules and 42 million spectra, which leverages a hybrid flow matching framework to orchestrate optimal transport paths for spectral peaks and discrete probability flows for molecular tokens. Furthermore, to guarantee the physicochemical validity of the generated structures, we incorporate explicit rule-based constraints into the generative process, utilizing a token-level mass control mechanism that strictly enforces alignment with the precursor molecular weight. MolSpecFlow establishes a new state-of-the-art on the MassSpecGym benchmark, surpassing diffusion and retrieval baselines in *de novo* generation (Top-1 accuracy +35%), spectral simulation, and molecular retrieval, demonstrating the power of physics-grounded unified modeling.

## 1. Introduction

Liquid chromatography-tandem mass spectrometry (LC-MS/MS) serves as the cornerstone for environmental analysis (Vermeulen et al., 2020), drug discovery (Dettmer et al., 2007; Atanasov et al., 2021), and disease diagnosis (Banerjee, 2020). However, over 90% of collected spectra remain unannotated “dark matter” (Wang et al., 2016; Alseekh et al., 2021), as current analysis is strictly bound by the closed-world assumption of database retrieval (da Silva et al., 2015; Li et al., 2021). Illuminating this space demands a shift toward unified generative modeling, integrating inverse structure elucidation (Stravs et al., 2022) to propose novel candidates and forward spectral simulation (Allen et al., 2015) to act as a rigorous verifier. Establishing this closed-loop system, however, faces two fundamental barriers.

The first challenge is modality heterogeneity. Mass spectra are not generic signals but sparse peak collections constrained by atomic masses (Qi & Volmer). Bridging the geometric gap between the discrete probability simplex of molecular graphs (Austin et al., 2021; Gat et al., 2024; Qin et al., 2024) and the physically constrained Euclidean manifold of spectra impedes synchronized generation (Lipman et al., 2022). Robust modeling must therefore reframe analysis from pattern matching to physics-grounded combinatorial optimization, where structural decisions are explicitly anchored by atom-level masses (Vaniya & Fiehn, 2015; Dührkop et al., 2019).

Second, a fundamental tension exists between scalable generation and structural representation. While permutation invariance aids representation learning (You et al., 2018; Kong et al., 2023), it creates computational bottlenecks for generative convergence (Vignac et al., 2022; Jo et al., 2022; Xu et al., 2024). Conversely, linearizing graphs (SMILES (Weininger, 1988)) enables scalability but introduces canonicalization bias (Bjerrum, 2017; Arús-Pous et al., 2019). Mirroring trends in protein design like AlphaFold 3 (Abramson et al., 2024), we posit that topological identity emerges not from strict equivariance, but as the consensus across the path group, relaxing constraints to favor generative fidelity.

To resolve these challenges, we introduce MolSpecFlow, a foundation model pre-trained on 100 million molecules (Irwin et al., 2020) and 42 million spectra (Bushuiev et al., 2025a). We propose a hybrid framework orchestrating discrete flow matching for molecular tokens and Optimal Transport paths (Liu et al., 2022) for spectral peaks. To navigate permutation sensitivity, we introduce a multi-path strategy that performs Monte Carlo integration over serialization orders, balancing representational robustness with generative flexibility. The overall framework is illustrated in Figure 4 (Appendix C), processing discrete tokens and continuous signals through a unified physics-aware embedding space.

Our main contributions are:

- **Unified Foundation Model at Scale:** We present the largest-scale multi-modal model to date, learning the joint distribution of molecular graphs and spectra to seamlessly support *de novo* generation, simulation, and retrieval.
- **Physics-Aware Hybrid Dynamics:** We formulate a Hybrid Flow Matching objective augmented by Token Mass Encoding that aligns discrete combinatorial flows with continuous physical signals.
- **Permutation Sensitivity Trade-off:** We investigate the intrinsic conflict between generation and representation. By modulating the number of paths, we reveal a tunable mechanism to balance structural consistency against decoding efficiency.

## 2. Related Work

### Physics-Informed Learning for Spectroscopy

AI has fundamentally transformed spectroscopy by prioritizing physics-informed representation learning (Guo et al., 2025). In NMR, methodologies have shifted from 2D graphs to 3D geometric deep learning for DFT-level prediction accuracy (Gerrard et al., 2020; Bhadauria et al., 2025), supporting inverse sequence generation tasks (Kim et al., 2023; Alberts et al., 2023). Similarly, advancements in IR/Raman and UV-Vis spectroscopy now incorporate equivariant networks to predict tensorial properties and excitation energies from quantum databases (Gastegger et al., 2017; Schütt et al., 2021; Singh et al., 2022; McNaughton et al., 2023). These approaches illustrate a paradigm shift toward unified foundation models capable of multimodal spectral reasoning (Guo et al., 2024).

### Generative and Representation Learning for Mass Spectrometry Analysis

Unlike non-destructive spectroscopy, MS involves complex fragmentation, prompting a shift from library searching (Aron et al., 2020; Lemmon et al., 2010) to learning continuous semantic embeddings. Early probabilistic representations (Huber et al., 2021a;b; Ortega et al., 2025) have evolved into sophisticated frameworks incorporating neutral loss encoding (Goldman et al., 2023a;b; Bui-Thi et al., 2024) and self-supervised pre-training on large-scale datasets (Bushuiev et al., 2025a;b; 2024). Parallelly, *de novo* generation (spec→mol) progressed from sequence translation (Shrivastava et al., 2021; Litsa et al., 2023; Butler et al., 2023) and two-stage pipelines (Stravs et al., 2022; Le et al., 2020; Neo et al., 2025) to topologically valid graph generation (Wang et al., 2025; Bohde et al., 2025). While forward simulation (mol→spec) has advanced via GNNs (Wang et al., 2021; Young et al., 2024; Overstreet et al., 2024), most approaches remain unidirectional, failing to close the generation-simulation loop required to resolve chemical ambiguity.

### Unified Molecular Generative Modeling under Complex Constraints

Current modeling efforts aim to integrate topological priors with physical constraints. Early SDE-based methods loosely coupled topology with noise but suffered from slow convergence (Huang et al., 2023; Zhang et al., 2023; Qiao et al., 2024; Feng et al., 2025). Alternatively, Molecular LLMs offer synthesis control but are limited by O(N) latency and ad-hoc handling of continuous modalities (Bagal et al., 2021; Zhang et al., 2024; 2025; Gruver et al., 2024). Flow Matching (FM) offers a potent resolution via deterministic Optimal Transport paths (Gat et al., 2024; Austin et al., 2021). Building on discrete FM explorations (Lee et al., 2025), we introduce MolSpecFlow, a unified framework synergizing discrete and continuous flows to combine linguistic precision with physical grounding for superior bidirectional generation.

## 3. Methods

We formulate mass spectrometry analysis as a generative modeling problem over the joint distribution of molecular structures and mass spectra. Unlike prior works that train separate models for forward (*x* → *s*) and inverse (*s* → *x*) prediction, MolSpecFlow learns a unified probability density *p*(𝒢, 𝒮) via a hybrid flow matching framework. This section establishes the problem definition, our unified pre-training strategy, the theoretical framework of path-marginalized molecular learning, and the hybrid flow matching objective.

### 3.1. Problem Formulation and Unified Modeling

#### 3.1.1. Notations and Definitions

Let 𝒢 denote the discrete space of molecular graphs. We represent a molecule as a sequence of tokens **x** ∈ 𝒳= {1, …, *V*} ^*L*^ (e.g., SMILES), where *V* is the vocabulary size and *L* is the sequence length. Let 𝒮 ⊂ ℝ^*N ×*2^ denote the continuous manifold of mass spectra, where each spectrum **s** ∈ 𝒮 consists of a set of peaks 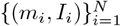 representing mass-to-charge ratios and intensities.

Our goal is to learn the joint distribution *p*(**x, s**). We decompose this objective into learning a time-dependent probability path *p*_*t*_(**z**) over the joint state space *Ƶ* = 𝒳 × 𝒮, evolving from a tractable prior distribution *p*_0_(**z**) = *p*_0_(**x**) × *p*_0_(**s**) to the data distribution *p*_1_(**z**) = *p*_data_(**x, s**).

To parameterize the vector fields governing these probability paths, we employ a unified Transformer architecture. Continuous spectral peaks are projected via a Learnable Gaussian Basis (LGB) encoder to avoid discretization artifacts, subsequently interacting with molecular tokens through a stack of multi-stream fusion blocks equipped with cross-modal attention. We refer readers to Appendix C for the complete mathematical formulations of the physics-informed embeddings, relative *m/z* bias, and layer-wise update dynamics.

#### 3.1.2. Mask-Conditioned Joint Pre-training

A critical challenge in scientific AI is the scarcity of paired data relative to the abundance of unimodal data. To leverage both, we propose a *Mask-Conditioned Joint Pre-training* strategy. We introduce a “dummy” state ∅ for each modality to represent missing or unconditioned information. The training objective approximates the joint log-likelihood by maximizing a mixture of conditional objectives over paired dataset 𝒟_pair_, molecular dataset 𝒟_mol_, and spectral dataset 𝒟_spec_:

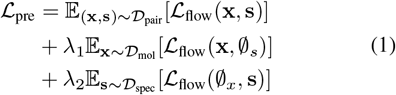

where ∅_*s*_ is a learnable embedding vector and ∅_*x*_ is a sequence of mask tokens. This formulation allows the model to simultaneously learn the joint correlation *p*(**x, s**) and the marginal distributions *p*(**x**) ≈ *p*(**x**|∅_*s*_) and *p*(**s**) ≈ *p*(**s**|∅_*x*_).

#### 3.1.3. Task Adaptation via Conditional Fine-tuning

Once pre-trained, the foundation model adapts to downstream tasks by modifying the conditioning context without architectural changes: (1) **Inverse Prediction (*de novo* Generation)** models the distribution *p*(**x**|**s**), where the spectrum **s** is clamped to *t* = 1 to guide the molecular flow **x**_*t*_ from noise to structure; (2) **Forward Prediction (Spectral Simulation)** models *p*(**s**|**x**), using the molecule **x** as the condition to generate spectral peaks **s**_*t*_; and (3) **Molecular Retrieval** models *p*(**h**_fp_|**s**), where the model is fine-tuned to predict molecular fingerprints **h**_fp_ (e.g., ECFP) from **s** to query large-scale databases via similarity ranking (e.g., Tanimoto coefficient).

### 3.2. Path-Marginalized Learning

#### 3.2.1. The Canonicalization Bias

A molecule is fundamentally a graph *G* ∈ 𝒢 invariant to node permutations. However, sequence-based models operate on linearizations (SMILES) *π* ∈ Π(*G*), where Π(*G*) is the set of valid traversals. Standard autoregressive models typically maximize the likelihood of a single canonical string *π*^∗^:

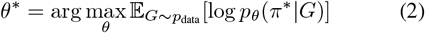

We argue this introduces a *Canonicalization Bias*: the model learns the grammar of the canonicalization algorithm rather than the topology of the molecule.

#### 3.2.2. Recovering Invariance via Multi-Path Integration

To approximate the true posterior *p*(*G*|𝒮), we must marginalize over the nuisance variable of graph traversal paths. We propose to learn the distribution over the path group Π(*G*) by maximizing the expectation:

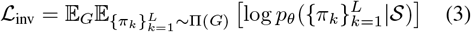

Unlike autoregressive models, our discrete flow matching framework operates globally. By processing a randomized ensemble of paths 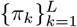 simultaneously and enabling information exchange via inter-path attention, MolSpecFlow performs Monte Carlo integration over the path space. The resulting representation, formed by averaging the outputs of the encoder Enc(·), approximates topological invariance:

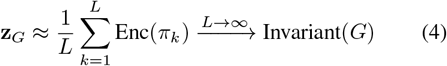

### 3.3. Hybrid Optimal Transport Flow Matching

We formulate the generative process as learning a time-dependent vector field that transports a prior distribution *p*_0_ to the data distribution *p*_1_. We propose a hybrid flow matching framework that orchestrates continuous dynamics for spectra/fingerprints and discrete dynamics for molecular tokens. Let *t* ∈ [0, 1] denote the flow time.

#### 3.3.1. Continuous Flow (Spectra and Fingerprints)

For continuous modalities **x**^cont^ (including spectral peaks **s** ∈ ℝ^*N ×*2^ and molecular fingerprints **h** ∈ ℝ^*D*^), we adopt the optimal transport conditional flow (Tong et al., 2023). The conditional probability path *p*_*t*_(**x**|**x**_1_) is defined as the push-forward of a Gaussian prior **x**_0_ ∼ 𝒩 (0, **I**) via a linear interpolant:

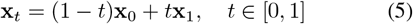

This path induces a constant target velocity field *u*_*t*_(**x**|**x**_1_) = **x**_1_ − **x**_0_. We train the neural velocity estimator 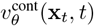 by minimizing the conditional flow matching regression loss:

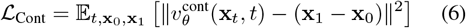

In our implementation, this objective is applied to both spectral peaks (ℒ_Spec_) and ECFP embeddings (ℒ_FP_).

#### 3.3.2. Discrete Flow on the Probability Simplex

For molecular tokens **x**^mol^ ∈ {1, …, *V*} ^*L*^, where *V* is the vocabulary size and *L* is the sequence length, we employ discrete flow matching on the probability simplex Δ^*V* −1^. Let *δ*_**x**_ denote the one-hot representation of token **x**. We define the conditional probability path *p*_*t*_(**x**|**x**_1_) as a mixture of the target distribution 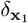 and a mask prior distribution 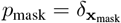, controlled by a scalar schedule *κ*(*t*):

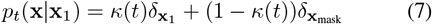

Here, *κ*(*t*) is a monotonically increasing function such that *κ*(0) = 0 and *κ*(1) = 1 (e.g., a cosine schedule). The training objective is to learn a probability denoiser *p*_*θ*_(**x**_1_|**x**_*t*_) that predicts the categorical distribution of the clean data **x**_1_ given the noisy state **x**_*t*_. The loss function is the expected cross-entropy over the sequence length *L*:

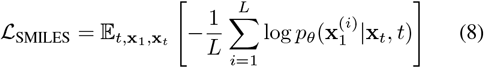

This objective guides the vector field to transport the probability mass from the mask token towards the correct token.

### 3.4. Physics-Constrained Objectives via Expectation

To ensure the generated structures adhere to physical laws, we impose constraints on the **expected** properties of the predicted distribution. Let **P** ∈ ℝ^*L×V*^ be the softmax probabilities predicted by the model at time *t*: 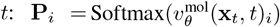.

#### Expected Formula Loss (ℒ_Formula_)

We map the token distribution to the elemental composition space using a constant registry matrix **C** ∈ ℝ^*V ×K*^, where *K* is the number of element types. The expected formula vector **ŷ** ∈ ℝ^*K*^ is computed via Einstein summation:

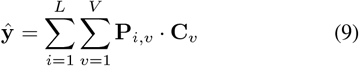

We minimize the Smooth L1 loss between **ŷ** and the ground truth formula **y**_gt_, applying a valid mask 𝕄_valid_ to handle dummy conditions:

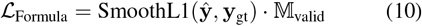

#### Expected Mass Loss (ℒ_Mass_)

Similarly, utilizing the atomic mass vector **w** ∈ ℝ^*V*^, we compute the expected total mass 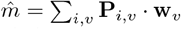. We constrain this expectation to match the precursor mass *m*_gt_:

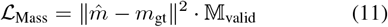

The total unified objective is a scalarized sum:

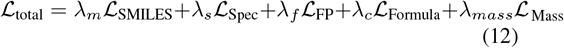

### 3.5. Coupled Inference via Synchronized Hybrid Euler Integration

Inference solves the joint Initial Value Problem (IVP) for the hybrid state 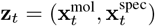 via a synchronized Euler scheme that evolves both manifolds simultaneously. At each step *t*, the model mutually predicts the spectral velocity 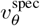 and molecular probabilities 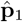. The continuous spectral state updates via a standard Euclidean step 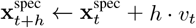. Simultaneously, for the discrete molecular sequence, we compute the probability vector field *u*_*t*_ on the simplex based on the flow balance equation (Gat et al., 2024):

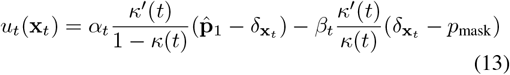

The next discrete state is then sampled stochastically from the evolved distribution: 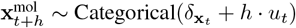. This process is governed by a cubic polynomial scheduler *κ*(*t*); detailed derivations of the update rules and the unified step size controller are provided in Appendix A and F.

## 4. Experiments

We evaluate MolSpecFlow on the MassSpecGym benchmark (Bushuiev et al., 2024), the largest and most rigorous standard for computational mass spectrometry. We focus on three core tasks: *de novo* molecular generation (inverse prediction), spectrum simulation (forward prediction), and molecular retrieval. Implementation details regarding datasets, architecture, and fine-tuning protocols are comprehensively provided in Appendices B, D, and G. Before quantitative analysis, we qualitatively validate the joint generation capability in Appendix H. Visualizations of complex structures exhibit a clear *coarse-to-fine* strategy: functional groups emerge early (*t* ≈ 0.15), core scaffolds stabilize by intermediate stages (*t* ≈ 0.45), and stereochemical details are refined in the final steps (*t >* 0.76). This evolution, perfectly synchronized with spectral convergence, confirms the model’s ability to effectively coordinate discrete molecular tokens with the continuous spectral manifold.

### 4.1. Experimental Setup

#### Dataset

We utilize the official MassSpecGym split, which comprises 231,104 high-quality tandem mass spectra (MS/MS) corresponding to 31,602 unique molecules. Crucially, this benchmark employs a Maximum Common Edge Subgraph (MCES) based splitting strategy. Unlike naive random splits or InChIKey-disjoint splits, the MCES split ensures that test molecules are structurally distinct from training molecules (edit distance ≥10), rigorously testing the model’s ability to generalize to ”dark chemical space” rather than simply memorizing spectral libraries.

#### Metrics

Following standard protocols (Bushuiev et al., 2024), we evaluate structure identification using Top-k Accuracy (exact match), MCES distance (structural edit distance), and Tanimoto Similarity (fingerprint overlap). For spectral simulation, we report Cosine and Jensen-Shannon similarities between binned spectral vectors. We strictly adhere to the MCES-based splitting protocol to ensure structural novelty. Due to space constraints, additional experimental results (e.g., retrieval tasks) and detailed metric definitions are provided in Appendix E.

### 4.2. De Novo Molecular Generation

We compare MolSpecFlow against state-of-the-art baselines including autoregressive sequence models (SMILES Transformer), retrieval-augmented generation (MADGEN (Wang et al., 2025)), and discrete diffusion models (DiffMS (Bohde et al., 2025)).

As shown in Table 1, MolSpecFlow establishes a new state-of-the-art. In the predictive setting (where no formula is provided), our model achieves a Top-1 Accuracy of 3.11%, surpassing the previous best, DiffMS (2.30%), by a relative margin of 35%. This improvement highlights the efficacy of hybrid flow matching in synchronizing the continuous spectral manifold with discrete molecular topology. Notably, the absolute Top-1 remains low, which is expected in MassSpecGym-style generalization splits: the spec→mol inverse problem is highly underdetermined, and exact-match Top-1 is dominated by search-space ambiguity rather than pure generative capacity.

**Table 1.** Performance comparison on MassSpecGym dataset. ”Ours” denotes the proposed MolSpecFlow model.

| Model | Top-1 |  |  | Top-10 |  |  |
| --- | --- | --- | --- | --- | --- | --- |
| | Accuracy ↑ | MCES ↓ | Tanimoto ↑ | Accuracy ↑ | MCES ↓ | Tanimoto ↑ |
| SMILES Transformer | 0.00% | 79.39 | 0.03 | 0.00% | 52.13 | 0.10 |
| MIST + MSNovelist | 0.00% | 45.55 | 0.06 | 0.00% | 30.13 | 0.15 |
| SELFIES Transformer | 0.00% | 38.88 | 0.08 | 0.00% | 26.87 | 0.13 |
| Spec2Mol | 0.00% | 37.76 | 0.12 | 0.00% | 29.40 | 0.16 |
| MIST + Neuraldecipher | 0.00% | 33.19 | 0.14 | 0.00% | 31.89 | 0.16 |
| Random Generation | 0.00% | 21.11 | 0.08 | 0.00% | 18.26 | 0.11 |
| MADGEN<sub>Pred.</sub> | 1.31% | 27.47 | 0.20 | 1.54% | 16.84 | 0.26 |
| MADGEN<sub>Oracle</sub> | 10.5% | 16.27 | 0.43 | 12.4% | 7.08 | 0.53 |
| DiffMS | 2.30% | 18.45 | 0.28 | 4.25% | 14.73 | 0.39 |
| MolSpecFlow<sub>pred</sub> | 3.11% | 17.32 | 0.34 | 5.23% | 12.32 | 0.44 |
| MolSpecFlow<sub>oracle</sub> | **13.6%** | **16.01** | **0.54** | **17.1%** | **6.95** | **0.60** |

Furthermore, our method demonstrates superior structural fidelity even when the exact molecule is not found. We achieve an MCES of 17.32 (lower is better) and a Tanimoto similarity of 0.34, significantly outperforming MADGEN (MCES 27.47) and DiffMS (MCES 18.45). This indicates that even incorrect predictions by MolSpecFlow conserve the core scaffolds and functional groups of the analyte more effectively than prior methods.

#### Oracle Performance

To disentangle the difficulty of formula prediction from structure generation, we evaluate an ”Oracle” setting where the ground-truth molecule scaffolds restricts the generation space (Wang et al., 2025). MolSpecFlow_*oracle*_ achieves 13.6% Top-1 accuracy, closely rivaling retrieval-based upper bounds and demonstrating that our generative backbone is highly effective when search space constraints are applied.

### 4.3. Spectrum Simulation and Retrieval Analysis

Beyond *de novo* generation, we evaluate MolSpecFlow as a bidirectional foundation model, assessing its capability to capture the probabilistic mapping between molecular topology and spectral fragmentation.

#### Forward Prediction

As shown in Table 2, MolSpecFlow achieves state-of-the-art performance in spectral simulation, recording a Cosine Similarity of 0.63 and a Jensen-Shannon (J.S.) Similarity of 0.59. Crucially, it significantly outperforms the specialized GNN-based baseline, FraGNNet (0.52 Cosine). We attribute this margin to two factors: (1) Modeling Long-range Dependencies: Traditional GNNs (like FraGNNet) are limited by message-passing depth, often failing to capture long-range rearrangement reactions typical in mass spectrometry (e.g., hydrogen scrambling). MolSpecFlow’s global attention mechanism effectively models these non-local interactions. (2) Continuous Intensity Modeling: The high J.S. similarity indicates that our Continuous Flow Matching objective accurately approximates the complex, multi-modal intensity distribution of ion fragments, avoiding the mode-collapse issues often seen in discrete binning approaches. The substantial improvement in Hit Rate (HR@1: 55.32% vs 46.64%) confirms that the model generates distinguishable spectral fingerprints unique to specific isomers.

**Table 2.** Performance comparison on spectral similarity and retrieval metrics. Values in parentheses denote confidence intervals. Bold indicates the best results.

| Model | Cosine Sim. ↑ | J.S. Sim. ↑ | HR @ 1 ↑ | HR @ 5 ↑ | HR @ 20 ↑ |
| --- | --- | --- | --- | --- | --- |
| Precursor m/z | 0.15 (0.14-0.17) | 0.15 (0.14-0.16) | 0.38 (0.21-0.62) | 1.72 (1.32-2.18) | 7.17 (6.32-8.04) |
| FFN Fingerprint | 0.25 (0.24-0.26) | 0.24 (0.23-0.25) | 8.44 (7.56-9.34) | 21.43 (20.10-22.79) | 38.57 (36.99-40.23) |
| GNN | 0.19 (0.18-0.20) | 0.20 (0.19-0.20) | 3.95 (3.37-4.62) | 11.92 (10.87-13.00) | 26.27 (24.83-27.82) |
| FraGNNet | 0.52 (0.51-0.53) | 0.47 (0.46-0.48) | 46.64 (44.98-48.26) | 72.56 (71.18-74.00) | 83.58 (82.34-84.75) |
| MolSpecFlow (Ours) | **0.63 (0.61-0.64)** | **0.59 (0.57-0.59)** | **55.32 (52.97-56.87)** | **79.98(79.01-81.32)** | **89.54 (88.32-90.76)** |

#### Molecular Retrieval

In the retrieval task (Table 3), MolSpecFlow acts as a neural ranker. Remarkably, our generative model surpasses MIST, the current state-of-the-art discriminative model trained specifically with contrastive learning (HR@1: 15.31% vs 14.64%). This result is nontrivial. Typically, specialized discriminative models outperform generative models on ranking tasks. MolSpecFlow’s superiority suggests that the joint flow matching pre-training learns a more chemically semantic latent space than pure contrastive loss. Furthermore, the MCES @ 1 (Maximum Common Edge Substructure distance of Top-1 predictions) drops to 14.32, significantly lower than MIST (15.37). This metric measures structural distance even when the exact match is missed. A lower MCES implies that even MolSpecFlow’s ”errors” are structurally homologous to the ground truth (Topological Smoothness), whereas baseline errors are often chemically irrelevant false positives. This property is critical for real-world discovery where the exact analyte may not exist in the database.

**Table 3.** Performance comparison on retrieval metrics and MCES. Values in parentheses denote confidence intervals. Bold indicates the best results.

| Model | Hit rate @ 1 ↑ | Hit rate @ 5 ↑ | Hit rate @ 20 ↑ | MCES @ 1 ↓ |
| --- | --- | --- | --- | --- |
| Random | 0.37 (0.24-0.54) | 2.01 (1.68-2.39) | 8.22 (7.53-8.89) | 30.81 (30.40-31.21) |
| DeepSets | 1.47 (1.18-1.77) | 6.21 (5.64-6.82) | 19.23 (18.24-20.26) | 25.11 (24.84-25.39) |
| Fingerprint FFN | 2.54 (2.17-2.99) | 7.59 (6.96-8.28) | 20.00 (19.01-20.98) | 24.66 (24.38-24.94) |
| DeepSets + Fourier features | 5.24 (4.71-5.83) | 12.58 (11.80-13.42) | 28.21 (27.10-29.36) | 22.13 (21.85-22.43) |
| MIST | 14.64 (13.82-15.54) | 34.87 (33.69-36.10) | 59.15 (57.89-60.39) | 15.37 (15.12-15.62) |
| MolSpecFlow (Ours) | **15.31 (14.45-16.21)** | **35.76(34.42-36.32)** | **61.32 (59.73-61.92)** | **14.32(14.15-14.98)** |

### 4.4. Scaling Laws and Path-Marginalized Dynamics

We investigate the scalability of MolSpecFlow along two distinct axes: pretraining scaling and inference compute (number of Monte Carlo paths).

#### Neural Scaling Laws and Emergent Capabilities

First, we analyze the impact of pretraining size by training variants ranging from 10M to 100M parameters. As illustrated in **Figure 2**, we observe a strict monotonic improvement in performance consistent with neural scaling laws. Notably, the performance gap widens significantly at the 100M scale (e.g., a *>* 4.5% absolute gain in HR@20 over smaller variants), suggesting that the complex probabilistic mapping between discrete molecular graphs and continuous spectra follows a non-saturating trajectory within this compute regime. This trend indicates that the mass spectrometry modality possesses sufficient information density to benefit from large-scale foundation models. The absence of performance plateaus suggests that further scaling will likely unlock emergent capabilities in distinguishing subtle stereoisomers and rare natural products, establishing MolSpecFlow as a scalable engine for future chemical foundation models.

**Figure 1.**
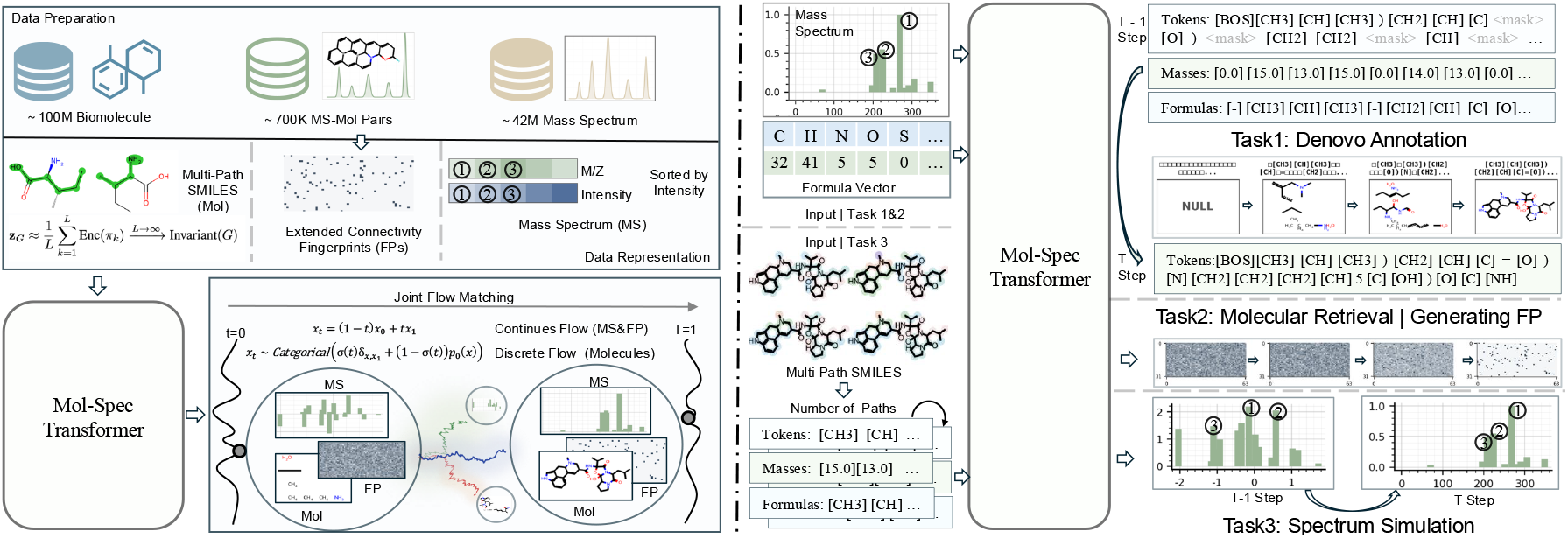
Overview of the MolSpecFlow Framework. (Top Left) Multi-Path Representation: Pre-trained on 100M molecules, the model aggregates randomized SMILES trajectories to approximate topological invariance, overcoming canonicalization bias. (Bottom Left) Joint Hybrid Flow Matching: The core unified Transformer learns to synchronize the evolution of diverse modalities. It couples discrete probability flows (for molecular tokens) with continuous optimal transport paths (for mass spectra and fingerprints), explicitly conditioned on chemical Formula Vectors. (Right) Physics-Aware Inference: The framework adapts to three downstream tasks: (1) *De Novo Annotation*, where generation is rigorously guided by token-level mass and formula constraints; (2) Molecular Retrieval via fingerprint prediction; and (3) Spectrum Simulation.

**Figure 2.**
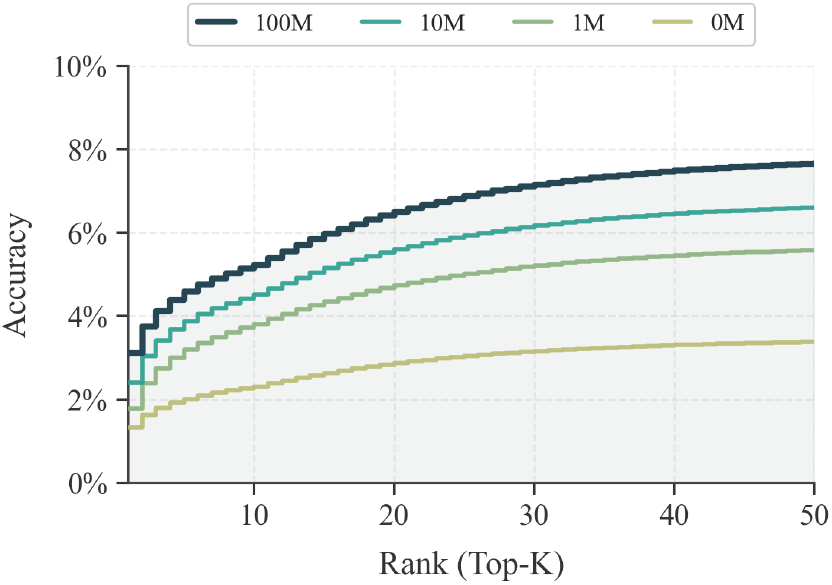
Neural scaling laws. Top-*K* accuracy improves monotonically as pretraining size increases (0M/1M/10M/100M).0M represents the baseline trained from scratch without pre-training.

#### Validating Path-Marginalized Sensitivity

Analyzing the impact of path number *L* reveals a striking divergence that empirically validates our theoretical dichotomy (Figure 3). In spectrum simulation (representation), performance (Hit@1) improves monotonically with *L*, confirming that Monte Carlo integration over randomized traversals effectively collapses the permutation space to recover a robust, invariant “consensus” embedding of the molecular topology. Conversely, in de novo generation, generative fidelity degrades as *L* increases. This phenomenon underscores the symmetry-scalability trade-off: while invariance aids representation, strictly enforcing the model to cover the high-entropy distribution of all possible SMILES permutations complicates the decoding process. The model struggles to collapse this combinatorial “superposition” of paths into a single syntactically valid sequence, highlighting the intrinsic computational burden of equivariant generation compared to the efficiency of invariant representation. Collectively, these results demonstrate that MolSpecFlow effectively resolves the symmetry-scalability trade-off: it leverages the scalability of Transformers (via scaling laws) while recovering graph invariance (via multi-path integration).

**Figure 3.**
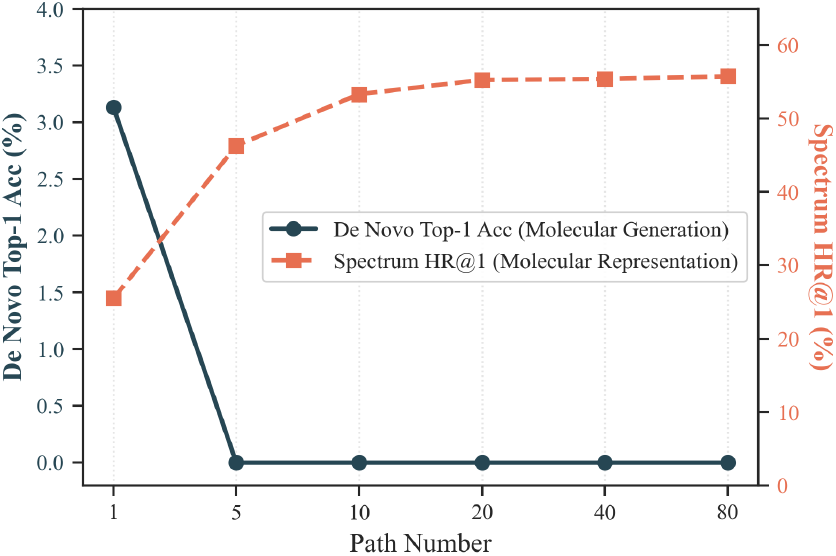
Path-marginalized dynamics. Increasing path number *L* improves Spectrum HR@1 (molecular representation) but degrades de novo Top-1 accuracy (molecular generation), revealing a symmetry–scalability trade-off.

**Figure 4.**
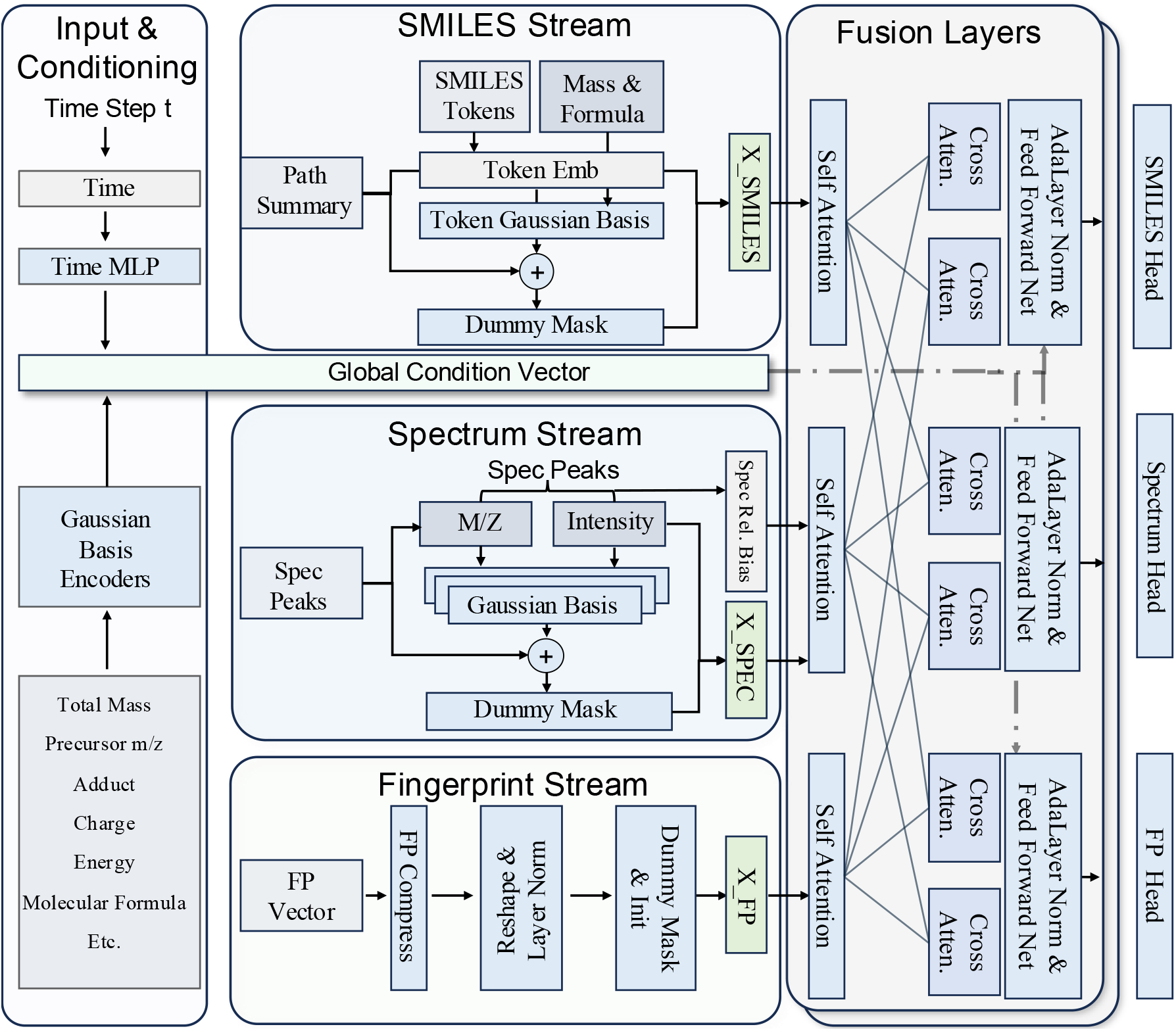
Detailed Architecture of the MolSpecFlow Transformer. The model features a three-stream structure processing **SMILES tokens** (Top), **Spectral Peaks** (Middle), and **Fingerprints** (Bottom). Continuous inputs (Mass, m/z, Intensity) are projected via *Token/Gaussian Basis Encoders* to avoid discretization errors. These streams interact through stacked **Fusion Layers**, which combine intra-modal *Self-Attention* with inter-modal *Cross-Attention* mechanisms. **Dash-dotted lines** indicate the injection of the Global Condition Vector (encoding precursor physics like Total Mass, Charge, Energy) and time-step embeddings into the *Adaptive Layer Normalization (AdaLN)* blocks, ensuring physics-aware modulation at every depth.

### 4.5. Ablation Study

To validate our architectural choices, we perform a comprehensive ablation study on the 100M-parameter pre-trained model (Table 4). The results highlight the criticality of continuous mass modeling: removing the Gaussian Kernel Projection leads to a 31% drop in Top-1 accuracy (3.11% → 2.15%), confirming that treating masses as continuous values via flow matching captures fine-grained precision far better than standard binning strategies. Furthermore, explicit constraints prove essential for structural validity; removing the deterministic formula embedding causes the MCES error to spike significantly from 17.32 to 23.80, suggesting that without conditional guidance, the discrete flow struggles to navigate towards chemically valid regions of the simplex. We also observe that the model relies heavily on local spectral context, as removing relative m/z positional encodings degrades Top-1 accuracy to 2.68%, hampering the ability to capture peak-to-peak distances and isotope patterns.

**Table 4.** Ablation study of key components on MassSpecGym dataset. All variants are based on the 100M pre-trained model.

| Model Variant | Top-1 |  |  | Top-10 |  |  |
| --- | --- | --- | --- | --- | --- | --- |
| | Accuracy ↑ | MCES ↓ | Tanimoto ↑ | Accuracy ↑ | MCES ↓ | Tanimoto ↑ |
| **Full Model (100M)** | **3.11 %** | **17.32** | **0.34** | **5.23 %** | **12.32** | **0.44** |
| w/o Mass Embedding | 2.15% | 21.45 | 0.28 | 3.92% | 16.10 | 0.36 |
| w/o Formula Embedding | 1.85% | 23.80 | 0.26 | 3.45% | 18.55 | 0.32 |
| w/o Relative MZ Pos | 2.68% | 18.92 | 0.31 | 4.75% | 13.88 | 0.41 |
| w/o All Above | 1.25% | 28.15 | 0.19 | 2.10% | 20.45 | 0.27 |

Ultimately, these components function synergistically rather than in isolation. As shown in the baseline where all proposed modules are removed, performance drops drastically to 1.25% Top-1 accuracy. This gap demonstrates that MolSpecFlow’s superior performance stems from the holistic integration of continuous mass embeddings, formula constraints, and positional awareness within the unified flow matching framework.

## 5. Conclusion and Future Work

In this work, we introduced MolSpecFlow, a unified foundation model that bridges discrete molecular graphs and continuous mass spectral signals via a physics-aware hybrid flow matching framework. A key factor is our multi-path training that couples spec→mol generation with mol→spec simulation and retrieval-style supervision, encouraging a shared representation that is both spectrum-consistent and structure-faithful, and improving robustness under experimental variability. Moreover, our token-level mass constraints act as an explicit physics prior that prunes mass-infeasible decoding trajectories early, effectively shrinking the search space and reducing chemically implausible “hallucinations”. This constraint is lightweight yet broadly applicable, leading to more reliable scaffold preservation (reflected by MCES/Tanimoto) even when exact-match identification is fundamentally ambiguous.

### Limitations and Discussion

While our results are promising, the ambiguity of the chemical space presents fundamental challenges for evaluation. A core limitation lies not in the generative capability, but in the criteria for correctness. The mapping between molecules and mass spectra is inherently many-to-many: distinct isomers can produce indistinguishable fragmentation patterns (spectral degeneracy), and a single molecule can yield variable spectra under different experimental conditions. Consequently, standard metrics like exact match accuracy or cosine similarity may penalize chemically plausible candidates that differ only in nuanced structural details from the ground truth. Future work must move beyond these rigid metrics toward physics-based evaluation, such as thermodynamic stability verification or consensus scoring across multiple instrument parameters. Additionally, extending MolSpecFlow to incorporate separation time (retention time) prediction could further constrain the search space, reducing ambiguity and paving the way for a truly comprehensive identification engine for the chemical “dark matter.”

### Software and Data

We will release the complete codebase (training/inference/evaluation), configurations, and pretrained/finetuned checkpoints upon acceptance. Molecular pretraining uses a 100M subset of ZINC22; spectrum-side pretraining uses the highquality filtered spectra subset DreaMS-A (as defined in GeMS/DreaMS); and benchmark experiments follow the official MassSpecGym benchmark datasets, splits, and metrics.

### Impact Statement

We develop physics-grounded ML models for MS/MS spectra to enhance chemical identification and structure elucidation. By improving analysis reliability and reducing experimental waste, our work offers significant benefits to metabolomics without notable negative impacts.

## A. Theoretical Derivations and Algorithms

### A.1. Discrete Flow Matching on the Simplex

We define the probability path *p*_*t*_(**x**|**x**_1_) as a linear interpolation between the target distribution 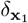 and a source mask distribution 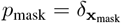. We define the vector field *u*_*t*_ to satisfy the continuity equation. The specific update rule employed in our framework is derived as:

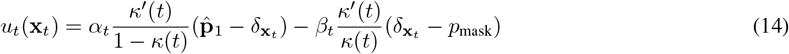

where 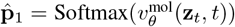 represents the predicted probability distribution over the clean data tokens. We utilize a scheduler *κ*(*t*) to control the flow dynamics.

### A.2. Unified Step Size Controller

To synchronize the dynamics of the discrete simplex flow and the continuous Euclidean flow, we implement a unified step size controller. This mechanism ensures that the probability mass transfer does not exceed a stability threshold *ρ* within a single Euler step. The dynamic step size *h*_*t*_ is calculated as:

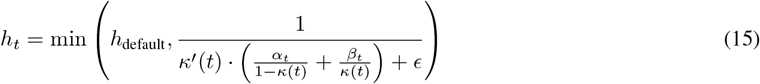

where *ϵ* is a small constant introduced for numerical stability.

### A.3. Inference Algorithm

The joint inference process employs a synchronized Euler integration scheme, optionally augmented with shallow fusion for the discrete modality to incorporate unconditional guidance. The complete procedure is outlined in Algorithm 1.

#### Algorithm 1

Synchronized Hybrid Euler Inference

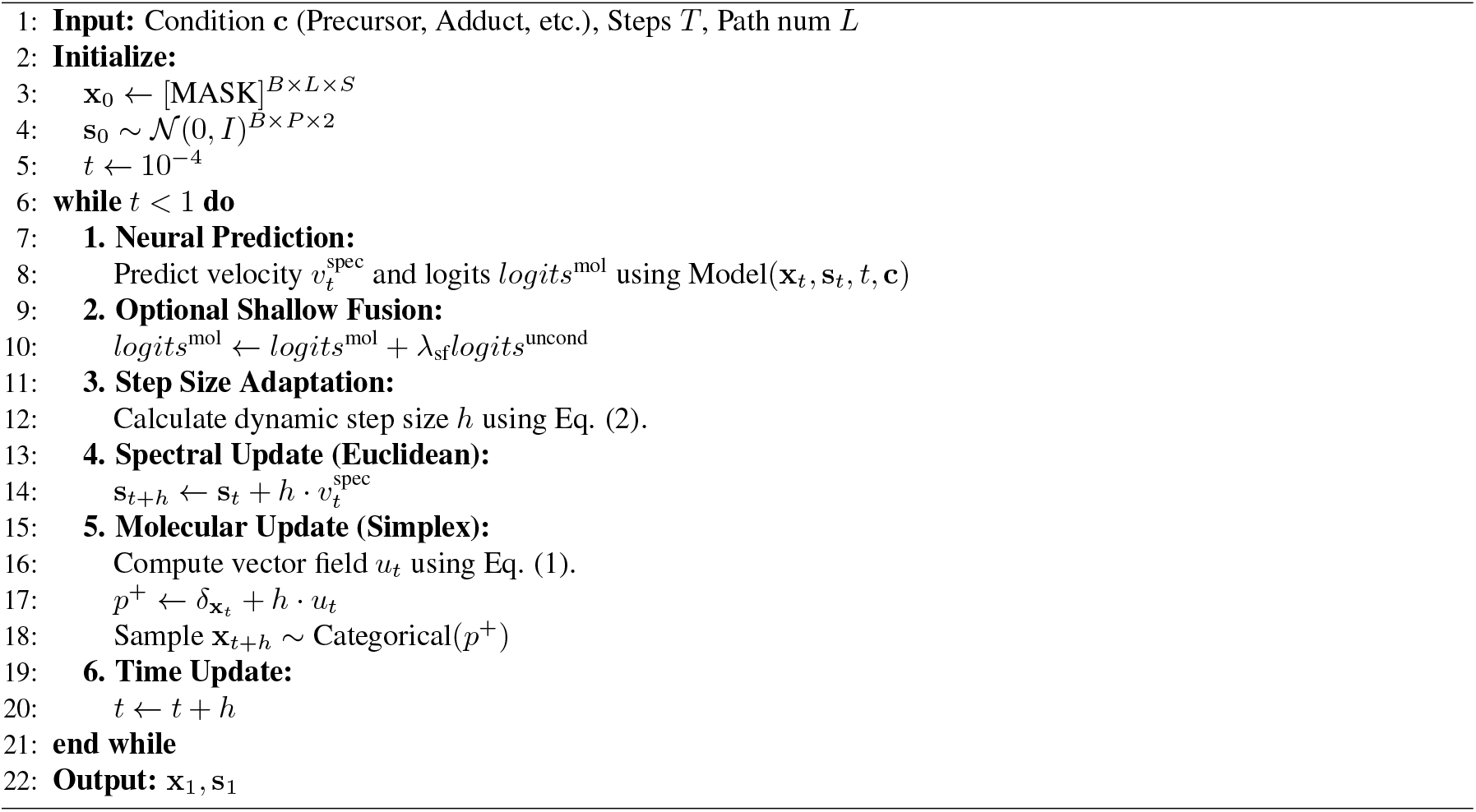

## B. Data and Preprocessing

### B.1. Datasets

We utilize three primary datasets:

- **ZINC 100M:** Used for large-scale pre-training of the molecular prior. It contains approximately 100 million drug-like molecules.
- **MassSpecGym:** A high-quality paired MS/MS dataset used for fine-tuning and evaluation. We strictly follow the MCES (Maximum Common Edge Subgraph) split to ensure structural separation between training and test sets.
- **GeMS:** A dataset of 42 million unlabeled mass spectra used to learn the continuous spectral manifold.

### B.2. Multi-Path Tokenization Strategy

To mitigate canonicalization bias and ensure physical precision, we employ a rigorous multi-path tokenization strategy:

#### 1. Explicit Hydrogen Materialization

Standard SMILES representations often rely on implicit hydrogen assumptions (e.g., ’c’ implies an aromatic carbon with one implicit hydrogen). However, this ambiguity is detrimental to physics-aware generation, as it complicates the precise calculation of mass and formula from discrete tokens. To resolve this, we strictly enforce **explicit hydrogen representation** (via RDKit’s `allHsExplicit=True`) before tokenization. This ensures that every hydrogen atom is materialized as a distinct token or explicit bracketed attribute. Consequently, the sum of the atomic masses of the tokens exactly equals the total molecular weight, enabling strict physicochemical validation without heuristic corrections.

#### 2. Randomized Path Augmentation

For each molecule *G*, we generate *L* distinct topological paths during training:

- The first path is the canonical DeepSMILES representation, serving as a stable reference.
- The subsequent *L* − 1 paths are generated by performing random graph traversals (using RDKit’s `doRandom=True`) followed by DeepSMILES encoding.

This strategy forces the model to learn a topological representation that is invariant to serialization order while strictly adhering to the mass conservation laws enforced by the explicit hydrogens.

#### 3. DeepSMILES Encoding

We utilize DeepSMILES (`rings=True, branches=True`) to handle ring closures and branching more robustly than standard SMILES, reducing the generation of syntactically invalid strings.

### B.3. Data Augmentation and Conditions

Input spectra are preprocessed to keep the top-200 peaks by intensity. For conditioning, we utilize:

- **Precursor** *m/z*: Normalized float value.
- **Adducts:** One-hot encoded or mapped to mass shifts (e.g., [*M* + *H*]^+^ → 1.007*Da*).
- **Collision Energy:** Normalized float value.

## C. Detailed Model Architecture

The MolSpec-Transformer processes discrete molecular tokens and continuous spectral signals through a unified physics-aware embedding space and a multi-stream fusion mechanism.

### C.1. Continuous Encoding via Learnable Gaussian Basis

To encode high-precision physical quantities (e.g., *m/z*, intensity, collision energy) without discretization artifacts, we employ a Learnable Gaussian Basis (LGB) encoder. For a scalar input *x* ∈ ℝ (e.g., peak intensity), the encoding 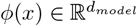 is computed as:

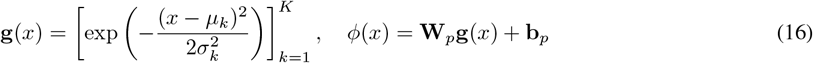

where 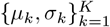 are learnable centers and widths initialized to cover the data range, and 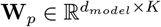 is a learnable projection matrix. This formulation allows the network to learn soft, overlapping receptive fields over the continuous physical manifold.

#### Algorithm 2

Physics-Informed Token Vocabulary Construction

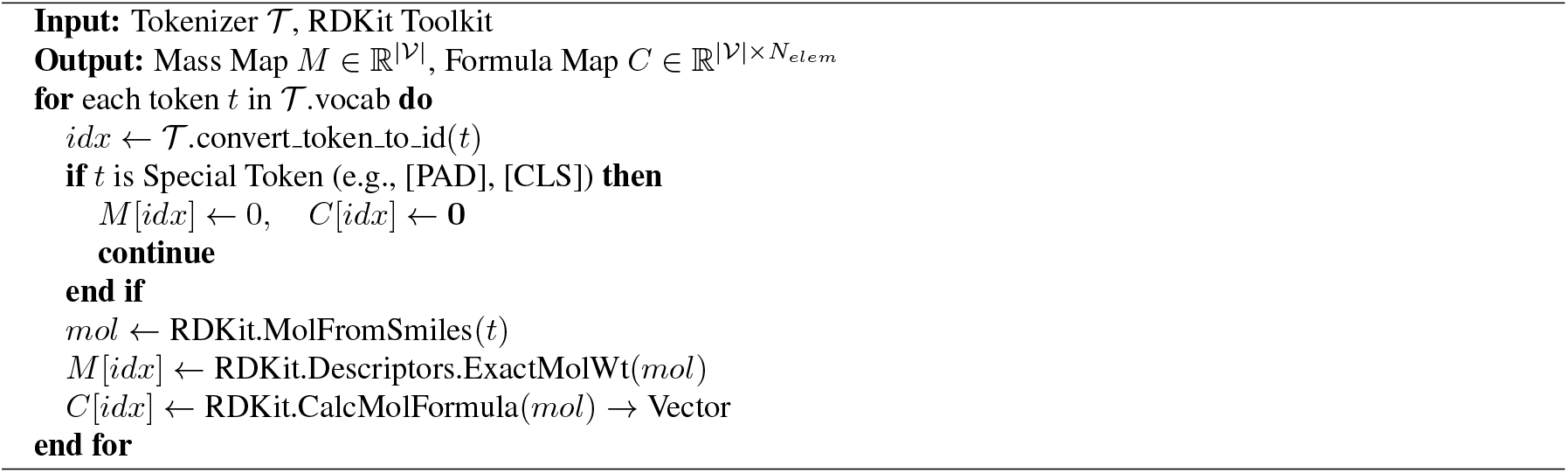

### C.2. Physics-Informed Token Embedding

Unlike standard language models that learn embeddings solely from distributional semantics, we enforce physical grounding by explicitly injecting mass and stoichiometric information into each token’s representation.

Let 𝒱 be the vocabulary. For each token *v* ∈ 𝒱, we pre-compute its exact monoisotopic mass *m*(*v*) and elemental composition vector 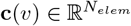 (Algorithm 2). The final embedding **h**_*i*_ for the *i*-th token *x*_*i*_ is a superposition of three components:

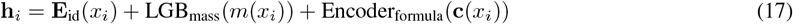

where **E**_id_ is a lookup table, LGB_mass_ is a Gaussian Basis encoder for mass, and Encoder_formula_ projects the sparse element count vector into the latent space.

### C.3. Relative m/z Bias Mechanism

In the spectral stream, to capture isotopic patterns and neutral losses invariant to absolute mass shifts, we introduce a **Relative m/z Bias** into the self-attention mechanism. For two peaks *i* and *j* with m/z values *z*_*i*_ and *z*_*j*_, the bias *B*_*ij*_ is computed as:

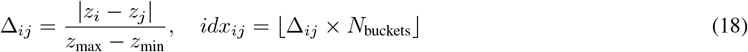

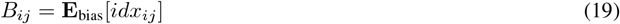

This bias is added to the attention scores: Attn_*ij*_ 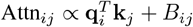.

### C.4. Multi-Stream Fusion Block

The model consists of *N* stacked fusion blocks that process the SMILES stream (**H**^mol^), Spectrum stream (**H**^spec^), and Fingerprint stream (**H**^fp^) in parallel. Global condition **c** (time *t*, precursor, etc.) modulates all layers via Adaptive Layer Normalization (AdaLN).

For layer *l*, the update dynamics are:

#### 1. Intra-Stream Processing (Self-Attention)

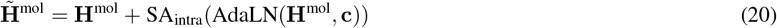

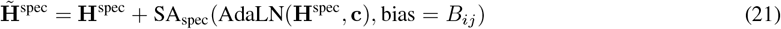

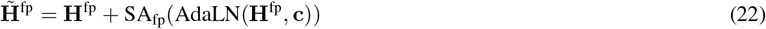

#### 2. Multi-Path Aggregation (SMILES only)

To enforce consistency across the *L* randomized SMILES paths, we perform inter-path attention using the path-averaged token representations:

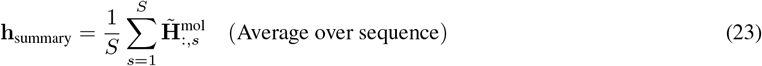

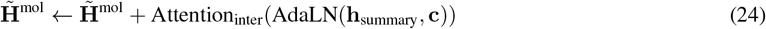

#### 3. Cross-Modal Fusion

Each stream queries information from the others. For the SMILES stream:

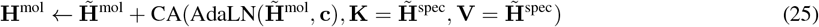

Similar cross-attention updates are applied to **H**^spec^ (attending to mol) and **H**^fp^.

#### 4. Feed-Forward

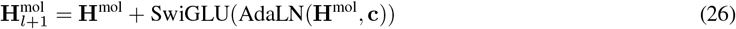

This architecture ensures that the discrete molecular generation is continuously guided by the spectral constraints and physical laws encoded in the conditioning vector.

## D. Training Formulation and Implementation Details

In this section, we present the comprehensive training protocol, including task formulation, detailed hyperparameters, and optimization settings derived from our codebase.

### D.1. Multi-Task Formulation

To enable robust bidirectional capability, we train the model on a mixture of three distinct objectives:

- **Joint Generation:** *x*_0_ = MASK, *s*_0_ ∼ 𝒩 (0, *I*). Both modalities are generated simultaneously from noise to model the joint distribution *p*(Mol, Spec).
- **Forward Prediction (Simulation):** *x*_*t*_ is fixed to the ground truth data. Only the spectral flow is optimized to predict spectra from molecules (*p*(Spec|Mol)).
- **Inverse Prediction (De Novo):** *s*_*t*_ is fixed to the ground truth data. Only the molecular discrete flow is optimized to predict molecules from spectra (*p*(Mol|Spec)).

### D.2. Model Configuration

Our model, MolSpec-Flow, is built upon a unified Transformer architecture. The specific hyperparameters are listed in Table 5.

**Table 5.**
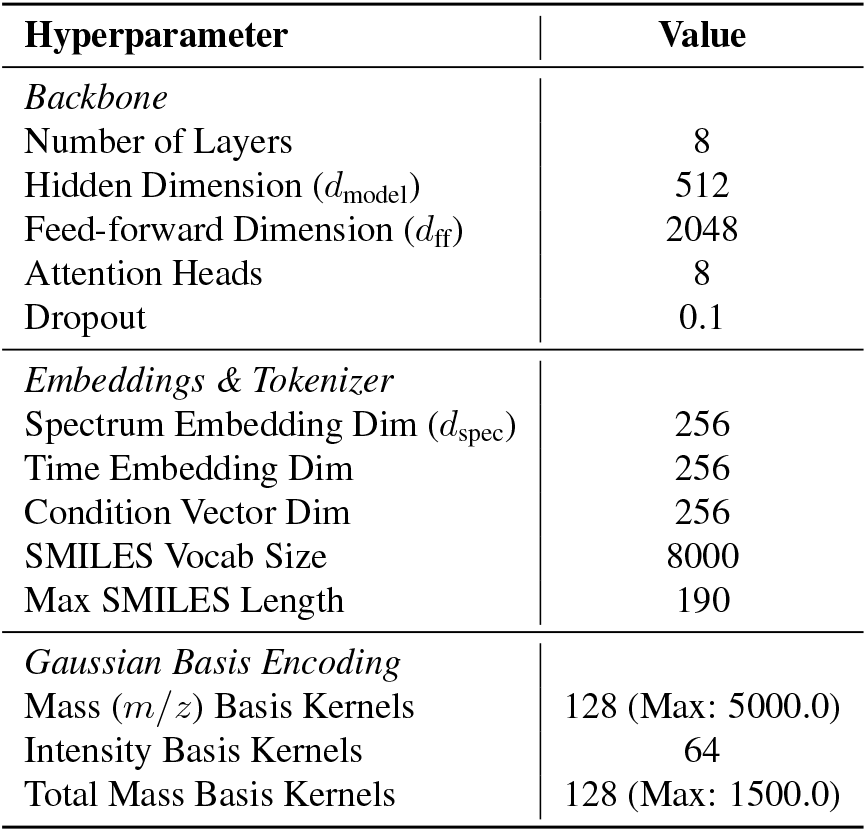
Detailed Hyperparameters of the MolSpec-Flow Architecture.

### D.3. Optimization Setup

We train the model using the **AdamW** optimizer with *β*_1_ = 0.9, *β*_2_ = 0.95, and a weight decay of 0.01. The global batch size is set to **320**. The learning rate follows a Cosine Annealing schedule with a peak value of 2 × 10^−4^ for the joint pre-training task. To stabilize the training and improve generation quality, we employ Exponential Moving Average (EMA) on model parameters with a decay rate of 0.999.

### D.4. Loss Balancing

The total training objective is a weighted sum of flow matching losses across different modalities. The specific weights used to balance these terms are:

- SMILES Reconstruction: *λ*_smiles_ = 1.0 (Pad token: 0.1)
- Spectrum Intensity: *λ*_intensity_ = 1.0
- Spectrum Position (*m/z*): *λ*_mz_ = 1 × 10^−5^
- Total Mass Prediction: *λ*_mass_ = 1 × 10^−6^

## E. Detailed Evaluation Protocols

We strictly adhere to the evaluation benchmarks and protocols established by MassSpecGym (Bushuiev et al., 2024) to ensure rigorous and reproducible comparisons.

### E.1. Dataset Splitting Strategy

To realistically simulate the challenge of identifying ”dark matter” in metabolomics, we utilize the **MCES-based Split** (Maximum Common Edge Subgraph). Unlike random splitting, which often leads to structural leakage (where isomers or stereoisomers of test molecules appear in the training set), the MCES split ensures structural disjointness.

- **Structural Independence:** The test set consists of molecules that are structurally distinct from the training set, defined by a strict MCES distance threshold.
- **Data Leakage Prevention:** This protocol guarantees that the model cannot simply memorize spectral fingerprints but must learn the generalized mapping between molecular topology and fragmentation patterns.

### E.2. Task 1: De Novo Molecular Generation (Inverse Prediction)

For the task of predicting a molecular graph *G* given a mass spectrum *s* (*p*(*G*|*s*)), we generate *K* candidates using beam search or sampling (typically *K* = 10). We evaluate performance using three complementary metrics:

#### 1. Top-*k* Accuracy (Exact Match)

The percentage of test samples where the ground truth molecule *G*_*gt*_ is strictly isomorphic to any of the top-*k* generated candidates *Ĝ*_1_, …, *Ĝ*_*k*_.

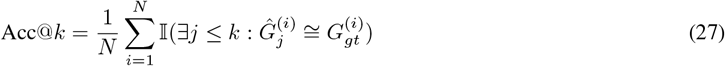

where ^∼^= denotes graph isomorphism checking (implemented via RDKit canonical SMILES matching).

#### 2. Tanimoto Similarity (Fingerprint Consistency)

To assess chemical similarity even when the exact match is missed, we compute the Tanimoto similarity between the Morgan fingerprints (Radius=2, 2048 bits) of the generated candidates and the ground truth.

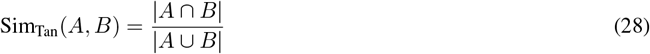

where *A* and *B* are the set of active bits in the fingerprint vectors. We report the maximum Tanimoto similarity among the top-*k* candidates.

#### 3. MCES Distance (Structural Distance)

The Maximum Common Edge Subgraph (MCES) distance quantifies the minimum number of edge edits (additions/deletions) required to transform the generated graph into the ground truth graph.

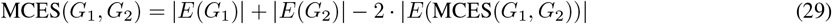

A lower MCES indicates higher structural preservation. This metric is robust to trivial functional group modifications that might drastically alter SMILES strings.

### E.3. Task 2: Spectral Simulation (Forward Prediction)

For the forward task *p*(*s*|*G*), we evaluate the similarity between the predicted spectrum *ŝ* and the ground truth spectrum *s*_*gt*_. Both spectra are binned (bin size = 0.1 Da) and normalized.

#### 1. Cosine Similarity

Measures the alignment of peak intensities in the high-dimensional spectral space:

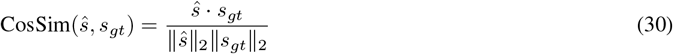

#### 2. Jensen-Shannon Similarity

To penalize discrepancies in peak distributions more rigorously than Cosine similarity (which is dominated by high-intensity peaks), we compute the Jensen-Shannon divergence (JSD) and report the similarity 1 − JSD.

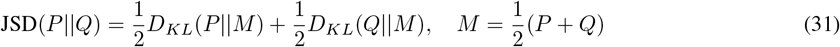

where *P* and *Q* are the probability distributions (normalized intensities) of the predicted and ground truth spectra.

### E.4. Task 3: Molecular Retrieval (Ranking)

In the retrieval setting, the model acts as a neural ranker to identify the correct molecule from a large candidate pool (e.g., PubChem) based on the query spectrum.

#### Protocol

Since MolSpecFlow is a generative model, we adapt it for retrieval by predicting the molecular fingerprint (ECFP) from the spectrum: *ĥ*_fp_ = *f*_*θ*_(*s*). We then rank all candidates in the library ℒ by calculating the Tanimoto similarity between the predicted fingerprint *ĥ*_fp_ and the candidate fingerprints *h*_*c*_.

#### Metric: Hit Rate @ *k* (HR@k)

The proportion of queries where the ground truth molecule appears in the top-*k* ranked candidates.

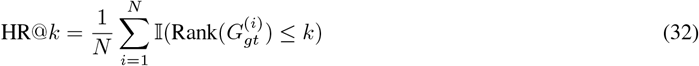

This setup specifically tests the model’s ability to map spectral features to the correct region of the chemical space, serving as a proxy for library-based identification performance.

## F. Generating Probability Velocity

The dynamics of the generative flow are governed by the scalar scheduler *κ*_*t*_, which dictates the rate of information transfer from the source mask distribution to the target data distribution.

To allow for flexible control over the flow velocity at the boundaries (*t* = 0 and *t* = 1), we adopt the parametric cubic polynomial scheduler proposed in prior work (Gat et al., 2024):

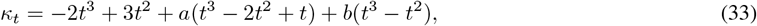

where *t* ∈ [0, 1], and *a, b* ∈ ℝ are hyperparameters controlling the initial and final slopes. This formulation strictly satisfies the boundary conditions *κ*_0_ = 0 and *κ*_1_ = 1. The derivative is given by:

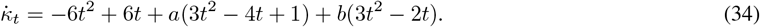

For our convex probability path, the marginal generating velocity field *u*_*t*_ for token *i* at state *z* is derived as:

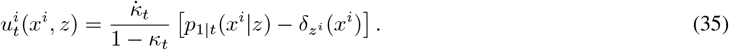

In our experiments, we set *a* = 2.0 and *b* = 0.5 to ensure a smooth transition profile that avoids singularities during sampling.

## G. Fine-tuning Protocol

To adapt the pre-trained foundation model to specific downstream tasks (e.g., *Spec-to-SMILES* or *Spec-to-ECFP*) while mitigating catastrophic forgetting, we employ a specific fine-tuning strategy composed of partial unfreezing, differential learning rates, and regularization.

### Layer-wise Unfreezing

Instead of fine-tuning all parameters, we freeze the majority of the backbone. We only unfreeze:

1. The task-specific projection heads (e.g., initialized linearly for ECFP or SMILES decoding).
2. The last 2 fusion layers of the Transformer backbone.
3. All LayerNorm parameters and biases, to allow feature distribution adaptation.

### Differential Learning Rates

We apply a lower learning rate to the pre-trained backbone layers to preserve general knowledge, and a higher learning rate to the task heads:

- **Backbone LR:** 2 × 10^−6^
- **Head LR:** 2 × 10^−5^

### L2-SP Regularization

For tasks with limited data, we utilize *L*^2^-SP (Starting Point) regularization. We add a penalty term 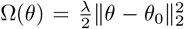, where *θ*_0_ are the pre-trained weights, constraining the fine-tuned model to stay close to the pre-trained manifold.

## H. Visualization of Generative Trajectories

In this section, we present the step-by-step generation process of MolSpecFlow. The figures illustrate the synchronized evolution of the molecular graph (Molecule Flow), the mass spectrum (Mass Spectrum Flow), and the molecular fingerprint (Fingerprint Flow) from noise (*t* = 0) to the final generated structure (*t* = 1).

**Sample 1:**
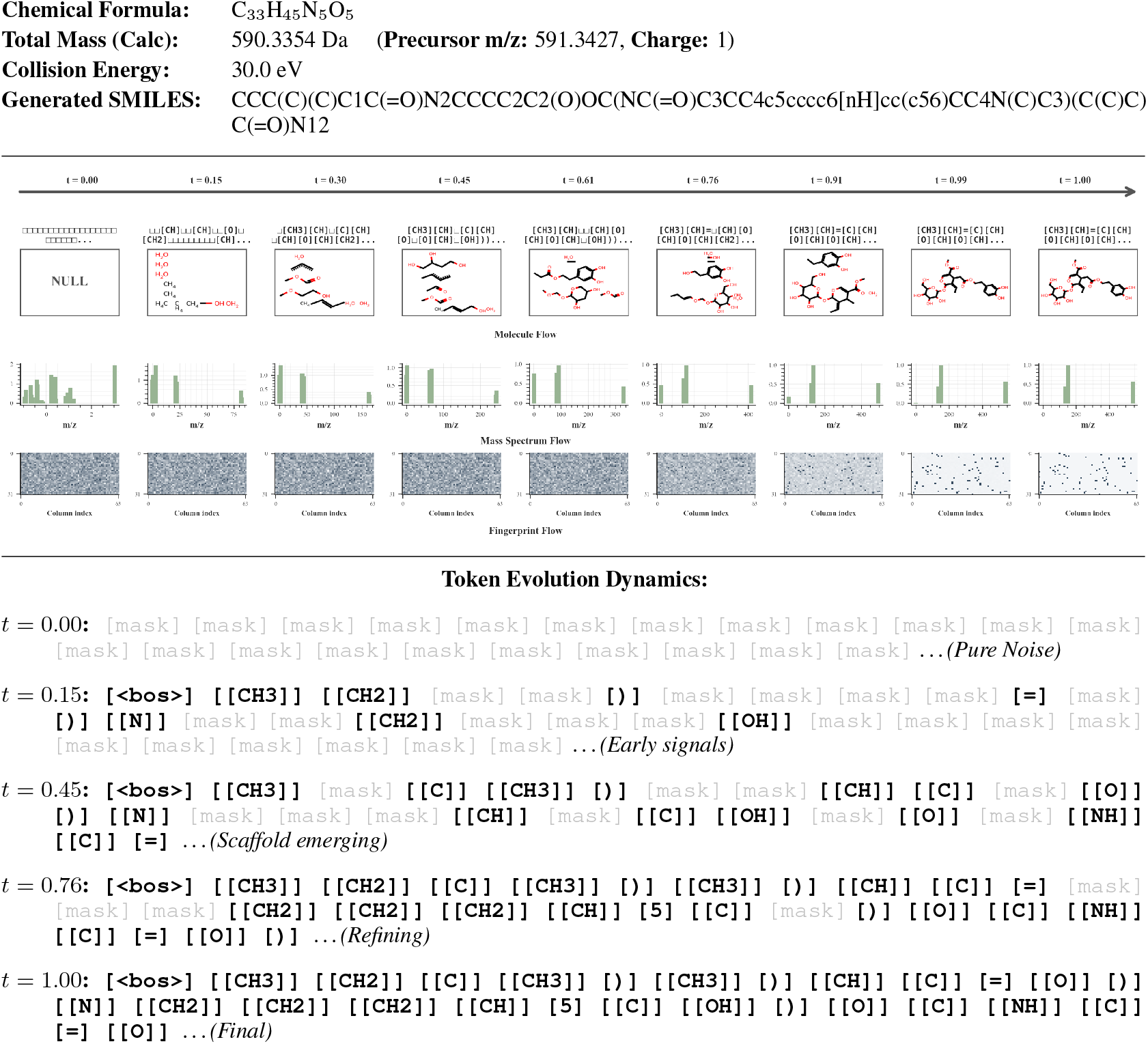
**Joint Generation of Peptidomimetic Structure**

**Sample 2:**
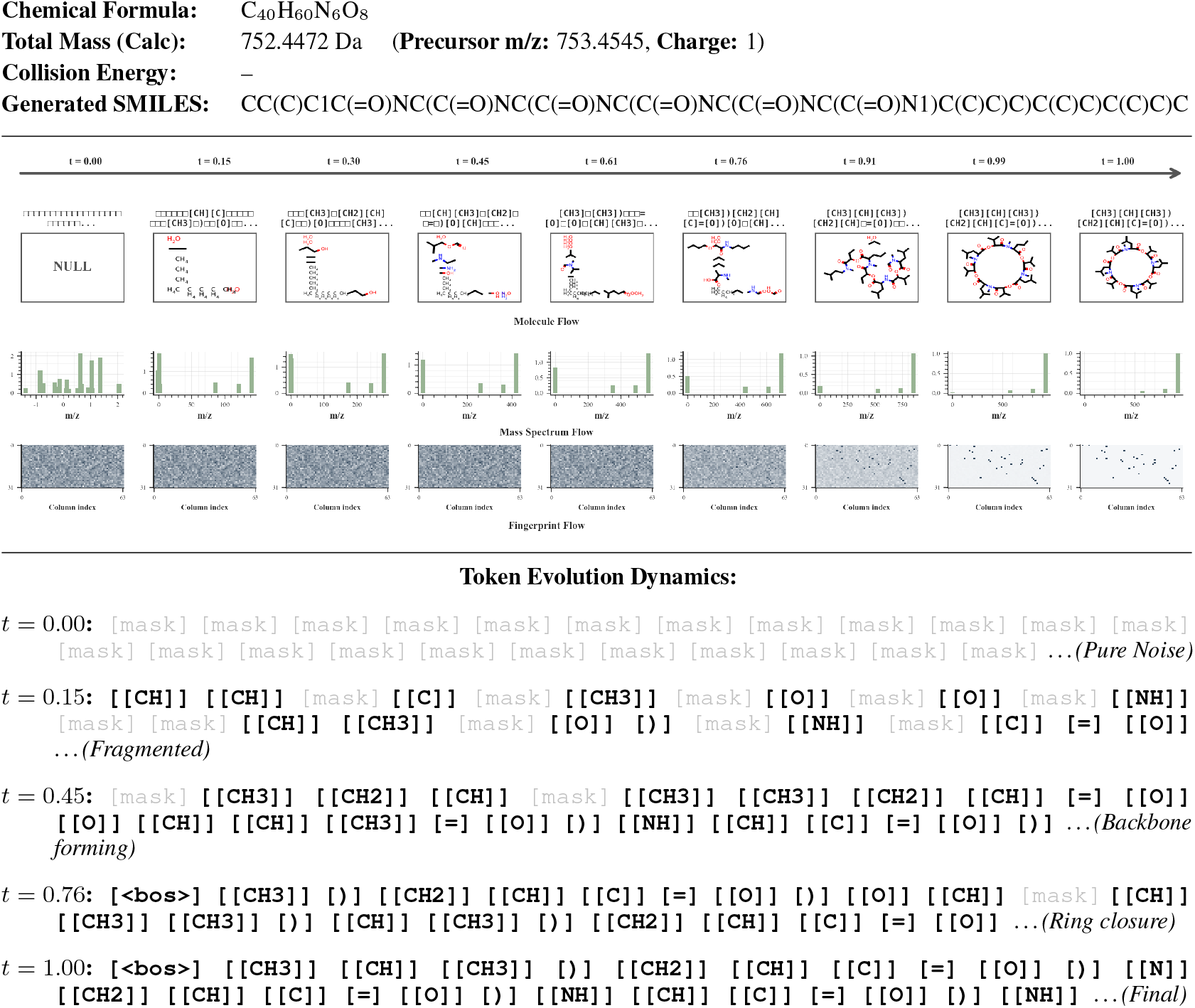
**Joint Generation of Macrocyclic Peptide**

**Sample 3:**
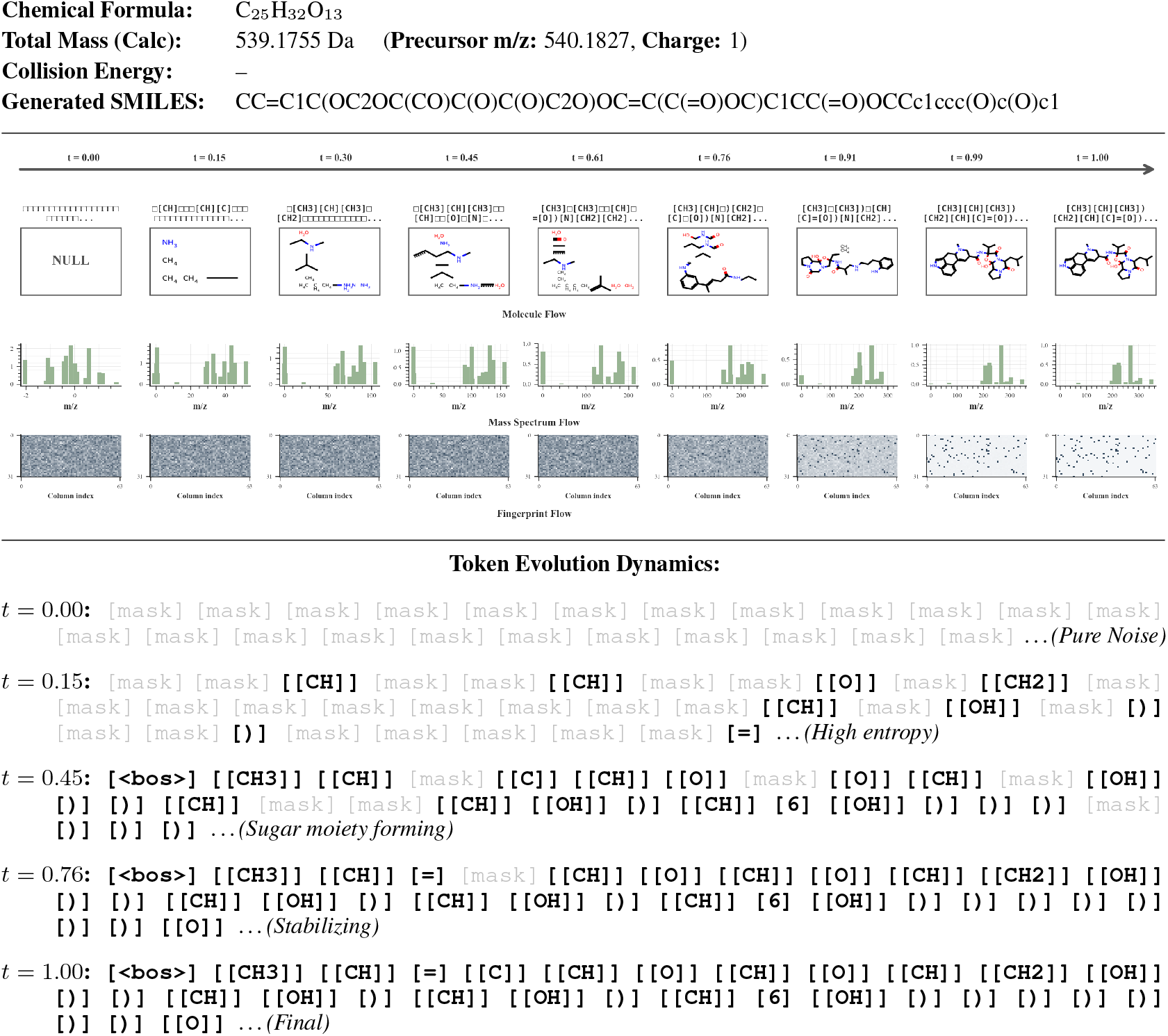
**Joint Generation of Glycoside Derivative**

